# *Plasmodium falciparum* invasion ligand gene transcript profiles in different populations

**DOI:** 10.64898/2026.04.15.718653

**Authors:** Elena Lantero-Escolar, Lindsay B. Stewart, Balbir Singh, Antoine Claessens, Alfred Amambua-Ngwa, David J. Conway

## Abstract

*Plasmodium falciparum* merozoites invade erythrocytes using various ligand-receptor interactions. Important ligands encoded by the *eba* and *Rh* gene families have varying expression levels in different parasite isolates, affecting their vaccine candidacy. Analyses of clinical isolates from endemic areas in Africa have indicated that most variation in these expression profiles exists within each local area, and only minor differences are seen between areas, although comparisons with non-African populations have not previously been performed. To enable this, relative transcript levels of three *eba* genes and five *Rh* genes have been analysed in new population samples, Malaysian isolates sampled from Sabah State in Borneo prior to endemic malaria elimination, and Gambian isolates, cultured under the same conditions to harvest schizonts for reverse transcription quantitative PCR assays. Significant differences between these populations were seen for three of the ligand genes, levels of *eba175* being higher in Malaysia, while levels of *eba181* and *Rh2b* were lower in Malaysia. The gene transcript profiles did not differ between single genotype and or multiple-genotype isolates. The distinctness of the Malaysian population expression profile was also supported by comparing previous data on clinical isolates from Ghana. In tests for correlation with previously determined parasite multiplication rates, *eba181* transcript levels correlated positively among Malaysian isolates but not among Gambian isolates. These findings suggest that expression of three *P. falciparum* merozoite ligands involved in invasion may be regionally differentiated, and further analysis of Asian parasite populations would be important if vaccines based on these candidates are to be considered for future use.

## 1. Introduction

Malaria remains a major global concern, and *P. falciparum* continues to be the causative agent of most malaria cases and deaths (World Health Organisation, 2024). The burden of *P. falciparum* infection in Africa remains very high, but Southeast Asian countries have made progress towards the long-term goal of eliminating this endemic parasite species. New tools are needed to achieve disease reduction and eventual elimination of malaria, and vaccines are likely to have a significant role (Ogwang and Crawley, 2025). The parasite mechanisms of erythrocyte invasion, and particularly the merozoite ligands involved, are candidate targets for vaccine development (Douglas et al., 2015; El Sahly et al., 2010; Payne et al., 2017) or possible treatment development (Srinivasan et al., 2013). Several Erythrocyte Binding Antigen (EBA) and Reticulocyte binding-like Homologue (Rh) proteins are encoded by genes transcribed at the parasite schizont stage, with products targeted to the apical organelles of merozoites where they function in erythrocyte invasion (Beeson et al. FEMS Microbiology Reviews 2016) (Lopaticki et al., 2011).

The expression pattern of the *eba* and *Rh* genes encoding these proteins varies between parasite clones cultured in the laboratory (Stubbs et al., 2005; Taylor et al., 2002) and also between different clinical isolates from which schizonts are studied in the first cycle of *ex vivo* development (Bowyer et al., 2015; Gomez-Escobar et al., 2010; Mensah-Brown et al., 2015; Nery et al., 2006). This expression is epigenetically regulated (Cortés et al., 2007), and changes occurring in culture can be selected (Jiang et al., 2010; Nyarko et al., 2020). Immune selection by antibodies against each of the individual ligands is considered a means of maintaining the range of variation of expression profiles *in vivo* (Persson et al., 2008). Studies on the variation in expression of merozoite ligand genes in clinical isolates have so far focused only on population samples from Africa. Most of the range of variation is seen within each local population studied, and there are only marginal differences between different populations (Bowyer et al., 2015; Mensah-Brown et al., 2015). However, *P. falciparum* populations outside of Africa remain understudied and may exhibit distinct patterns of variation.

In this study, we collected RNA samples after adaptation to culture of *P. falciparum* archived isolates from two different geographical locations: The Gambia and Malaysia Borneo. To our knowledge, this the first study to quantify invasion-ligand transcript levels in isolates of South East Asian origin using this approach. Our goal was to determine whether these isolates display distinct expression patterns compared with West African parasites. In addition, we hypothesised that transcript profiles of invasion ligands may correlate with PMRs, as both may be subject to similar selective pressures *in vivo*.

## 2. Methods

### 2.1. *P. falciparum* clinical isolates

Twenty-one Malaysian *P. falciparum* isolates were analysed, originally sampled from Sabah state in 1996 and 1997, with blood samples being cryopreserved and parasites cultured as recently described (Stewart et al., 2025b). Briefly, samples had been collected into EDTA blood collection tubes, with aliquots cryopreserved in liquid nitrogen in Malaysia and then sent on dry ice to the London School of Hygiene and Tropical Medicine (LSHTM) for continued storage in liquid nitrogen until thawing for parasite culture (Stewart et al., 2025b).

Twenty Gambian clinical isolates were also analysed, originally sampled from the Upper River Region of The Gambia in 2014 and 2016, with blood samples being cryopreserved and parasites cultured as recently described (Stewart et al., 2025a). Briefly, samples had been collected into heparinised vacutainer tubes, with aliquots cryopreserved in liquid nitrogen in The Gambia, and then sent on dry ice to LSHTM for continued storage in liquid nitrogen until thawing for parasite culture (Stewart et al., 2025a).

### 2.2. Parasite culture

Culture of these clinical isolates has been previously described in studies of *P. falciparum* multiplication rate variation (Stewart et al., 2025a, 2025b). Briefly, thawing from cryopreservation was performed, and *P. falciparum* parasites were cultured continuously at 37 °C using standard methods (Trager and Jensen, 1976), in RPMI 1640 medium supplemented with 0.5% Albumax II, under an atmosphere of 5% O_2_,5% CO_2_, and 90% N_2_, with orbital shaking of 6-well plates at 60 revolutions per minute. Isolates were cultured for at least two weeks, by which time most erythrocytes had been replaced by those from non-patient donors. Numbers of different parasite genotypes in genomic DNA from each of the isolates were previously assessed by genotyping the polymorphic *msp1* and *msp2* genes (Stewart et al., 2025a, 2025b), and these data were used here for comparing single and multiple-genotype isolates. For each isolate, the numbers of days in culture before harvesting of schizonts for gene transcript analysis are given in Supplementary Table S1.

### 2.3. RNA extraction from schizonts for transcript analysis

A total of 41 parasite isolates (21 from Malaysia and 20 from The Gambia) were analysed, out of 64 parasite isolates cultured in attempt to obtain schizont RNA, most of which had been previously included in analysis of parasite multiplication rates (Stewart et al., 2025a, 2025b). From each isolate, MACs purification of mature intra-erythrocytic parasites (pigmented trophozoite and schizont stages) was performed between days 11 and 35 of starting parasite culture, two or three-time points being collected and pooled to ensure enough material for analysis. Then, parasites at 0.8% haematocrit (HTC) (adjusted with some RBC) were allowed to mature for 2-5 hours in presence of 10 µM E-64, by which point segmented schizonts were the majority of the parasites observed, and these were collected for RNA extraction. Erythrocytes were then resuspended at 50% haematocrit and immediately mixed with 4 volumes of TRIzol reagent (Ambion). Aliquots were stored at -80 °C for subsequent RNA extraction using a RNeasy Micro kit (Qiagen) and quantified with Qubit RNA kit. Only samples with RNA concentration higher than 2.5 ng µL^-1^ were included in the subsequent analyses (41 isolates). The poly-adenylated mRNA was reverse transcribed with oligo(dT) using TaqMan reagents (Applied Biosystems). The resulting complementary DNA was used for real-time quantitative polymerase chain reaction (RT-qPCR) assay with gene specific TaqMan primers and probe sets using the same methods as previously applied in the same laboratory in analyses of schizonts from the first *ex vivo* generation in clinical isolates from different West African populations (Bowyer et al. 2015). The primer and probe sequences for each gene are as previously described elsewhere, for the *Rh1, Rh2a, Rh2b*, and *Rh4* genes (Nery et al., 2006), for *eba140, eba175*, and *eba181* (Blair et al., 2002), and for the *Rh5* (Gomez-Escobar et al., 2010). The RT-qPCR reactions were performed in 10 µL volumes with 900 nM of each primer and 250 nM of each probe, using the 7500 Real-Time PCR system (Applied Biosystems) with the first cycle at 50°C for 2 min and 95°C for 10 min, followed by 40 cycles of 95°C for 15 s and 60°C for 1 min. Each run included controls and 3D7 genomic DNA standards, with each sample assayed in duplicate and standard curves generated during each run.

### 2.4. *eba* and *Rh* gene expression analysis

The relative transcript level of each of the *Rh* and *eba* genes was derived as a normalized proportion of the sum of the transcript levels for the 8 genes *(eba140, eba175, eba181, Rh1, Rh2a, Rh2b, Rh4* and *Rh5)* in each isolate. Spearman’s rho was used to assess the nonparametric rank correlations, and Mann-Whitney U tests were used to assess whether there were significant differences between groups in the distributions of continuous variables. Only results with p < 0.01 were considered significant. Both Spearman’s rho and Mann-Whitney U tests were applied using GraphPad PRISM 10.

## 3. Results

### 3.1. Comparing *P. falciparum eba* and *Rh* gene expression profiles among clinical isolates from Malaysia and The Gambia

Across the 41 cultured clinical isolates from Malaysia (N = 21) and The Gambia (N = 20) the eight *eba* and *Rh* ligand displayed a broad range of expression levels, with *eba175* and *eba140* being the most predominant of these transcripts (Fig. 1). A summary of the varying profiles is illustrated as a heatmap (Fig. 1A) and the values for each of the isolates are tabulated (Supplementary Table S1). Although both populations exhibit substantial within-country variation, clear differences emerge when comparing relative transcript levels between populations. Notably, *eba175* has relatively higher expression in Malaysian isolates (p = 0.0019), while conversely *eba181* (p < 0.0001) and *Rh2b* (p < 0.0001) have relatively higher expression in Gambian isolates (Fig. 1B). No significant differences were seen between isolates that contained single parasite genotypes and those with multiple genotypes (Fig. 1C).

**Figure 1:**
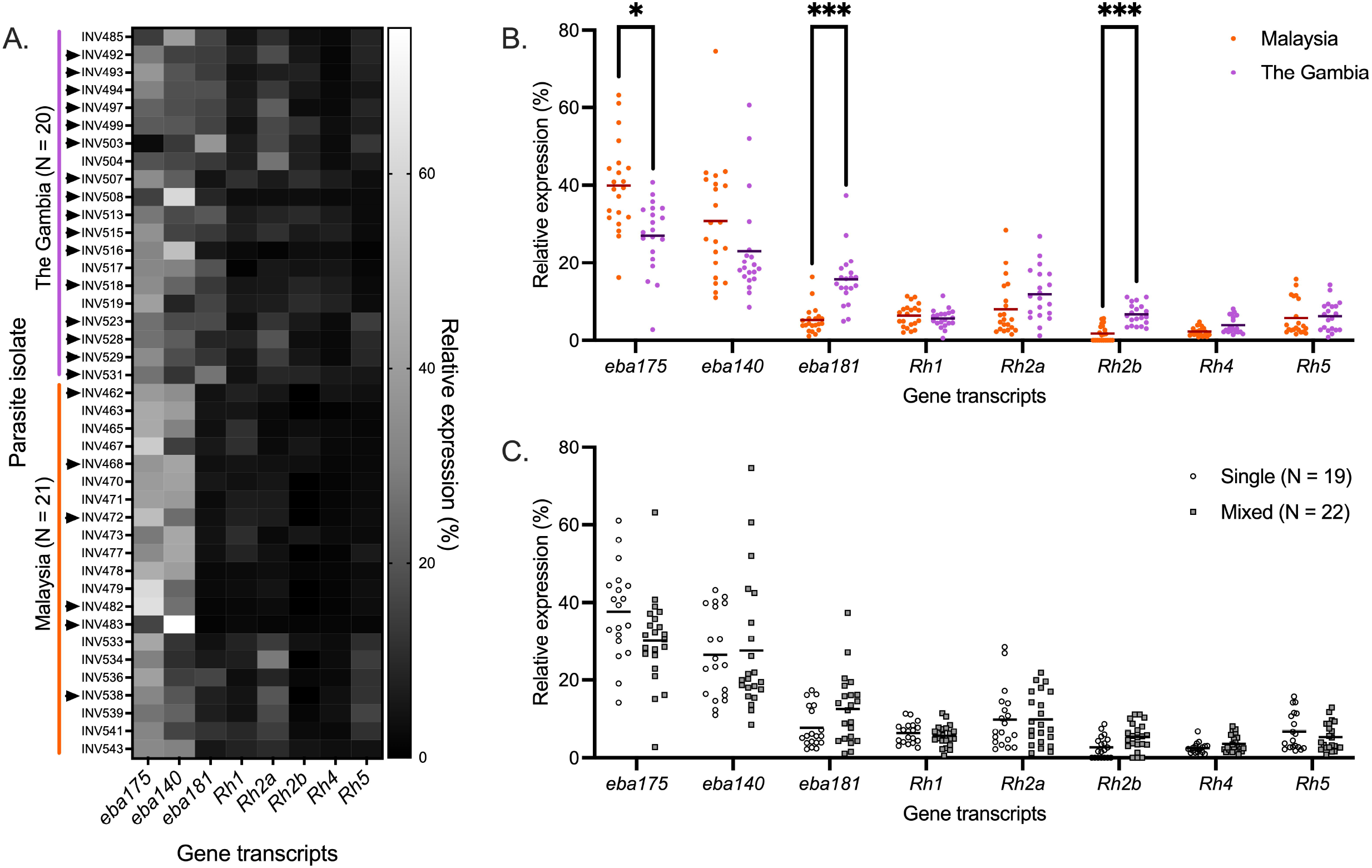
Relative expression levels of *eba* and *Rh* families expressed in percentage. **A:** Heatmap showing the levels of expression in each one of the lines assayed. Arrowheads indicate multiple genotypes present in the sample. **B:** Transcript levels grouped by origin, Malaysian Borneo samples show higher levels of *eba175* (P = 0.0019) while the expression of *eba181* (P < 0.0001) and *Rh2b* (P = 0.0001) is higher in Gambian samples. Horizontal lines represent the mean relative expression of the sample groups. **C:** Relative transcript levels grouped by the number of genotypes found in the samples, either single genotype or more than 2 genotypes (considered mixed); not significant differences can be observed due to presence of just one or two or more genotypes. Horizontal lines represent the mean relative expression of the sample groups. Statistical analysis with Mann-Whitney t-test in GraphPad PRISM 10. Significant values are labelled with * for P < 0.01, with ** for P < 0.001 and with *** for P < 0.0001.

**Figure 2:**
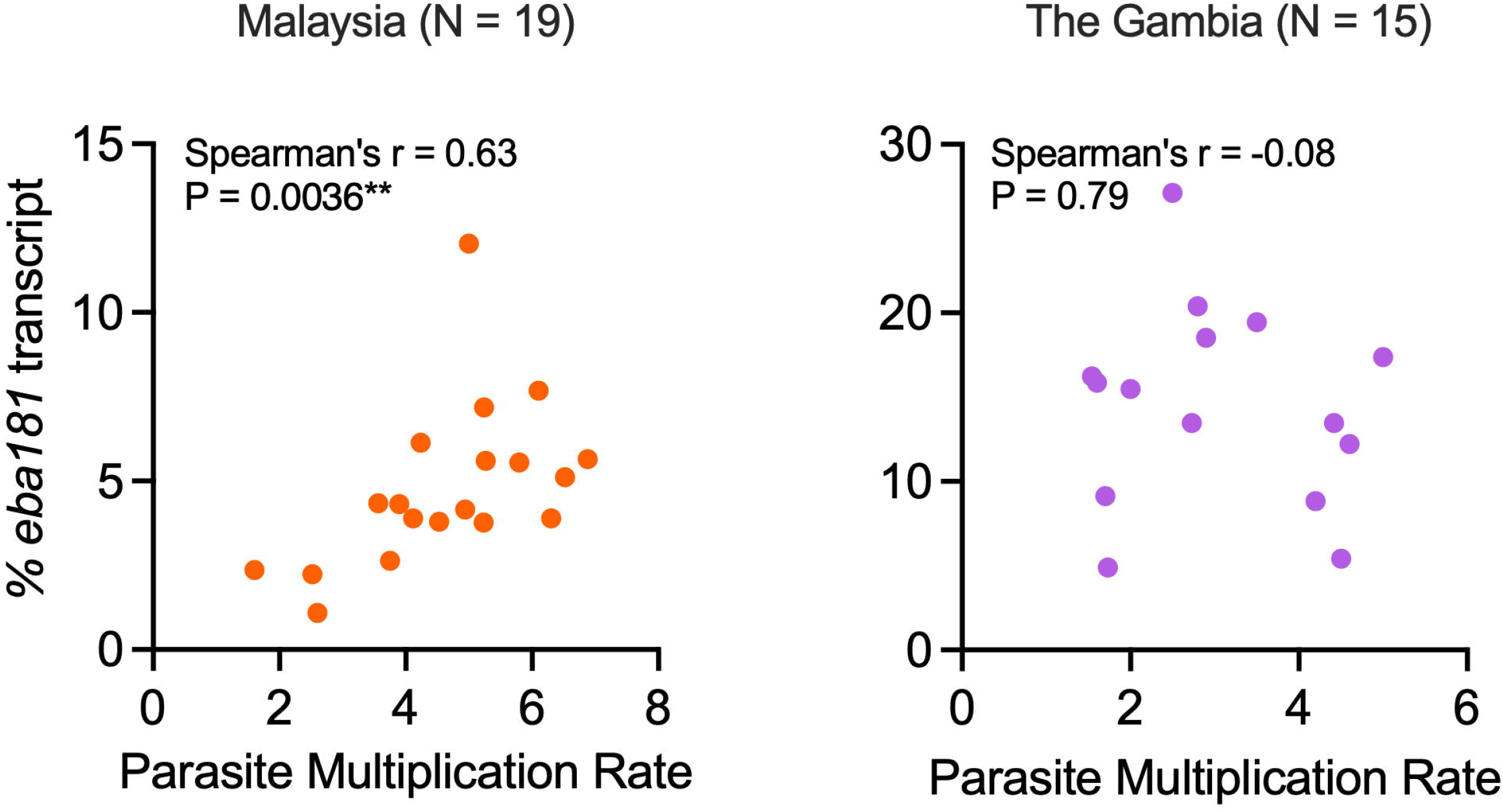
Scatterplots of *eba181* transcript ratios vs PMR of the assayed lines. Both populations assayed are plotted with distinct colours. % of *eba181* transcripts only show a significant correlation with PMR in the samples of Malaysian Borneo origin. Analysis in GraphPad PRISM 10.

### 3.2. Testing for correlations between transcript profiles and multiplication rates

*In vitro* exponential multiplication rates were previously measured for thirty-four of these *P. falciparum* isolates (19 Malaysian and 15 Gambian isolates) (Stewart et al., 2025a, 2025b) (Supplementary Table S1), enabling tests for correlations between merozoite ligand transcript variation and growth rates. For each population separately, nonparametric rank correlations were calculated between relative expression levels of the *eba* and *Rh* ligand genes and parasite multiplication rates (Table 1). Among the Malaysian isolates, *eba181* transcript levels in schizonts correlated positively with the parasite multiplication rates (Spearman’s rho = 0.63, P = 0.0036), but there was no significant correlation among the Gambian isolates (Fig. 3 and Table 1). None of the other gene transcripts showed any significant correlation with parasite multiplication rates (Table 1), and most of the non-significant trends in positive or negative directions are not reproduced in the tests on the different populations.

**Table 1:**
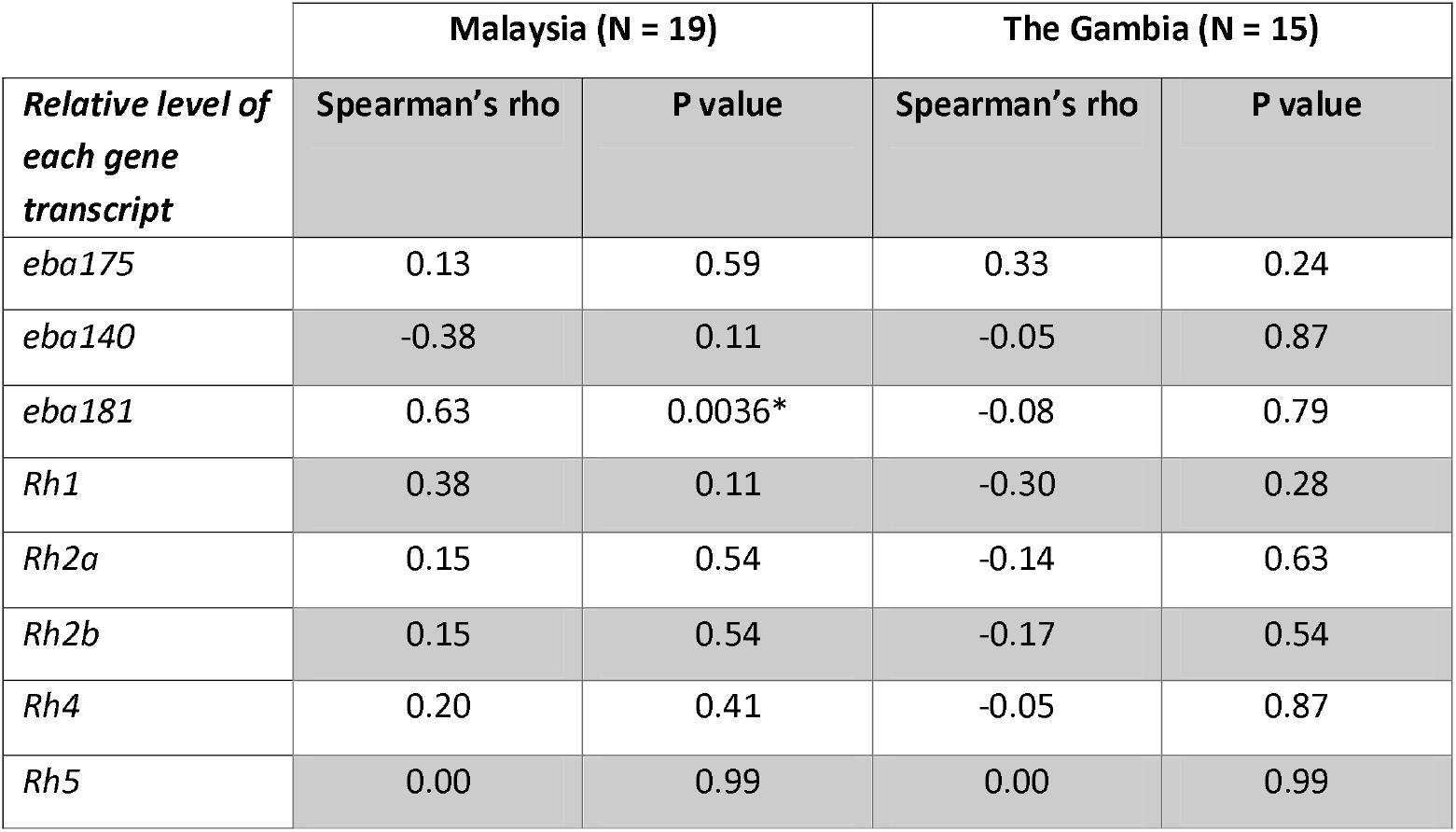
Tests for correlation between relative levels of individual *eba* and *Rh* gene transcripts in schizonts and parasite multiplication rates in Malaysian and Gambian *P. falciparum* isolates.

**Figure 3:**
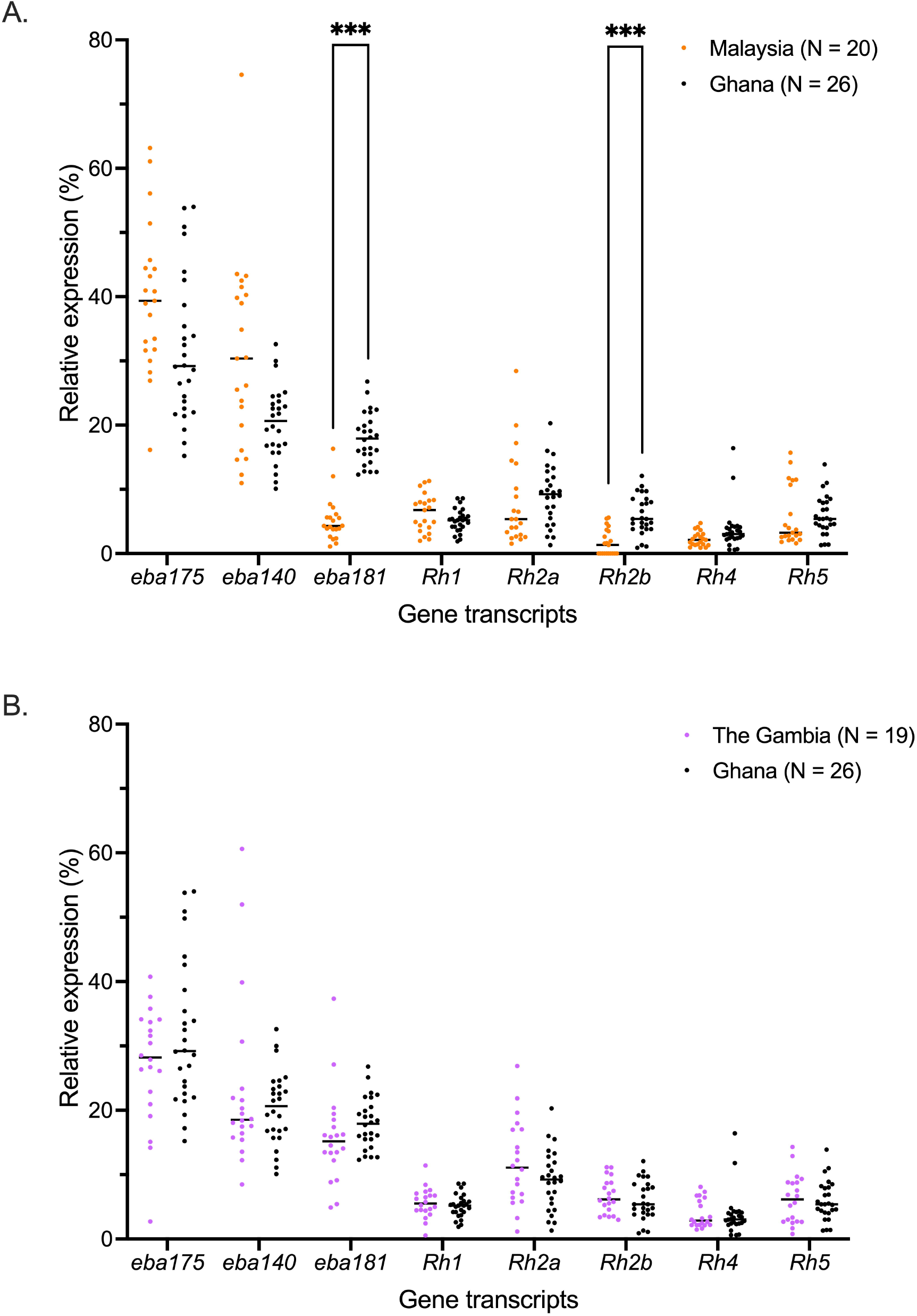
**Comparisons of the relative expression of the invasion ligands between Ghanian samples analysed in a previous study** (Bowyer et al., 2015) **and the Malaysian (A) and Gambian (B) samples analysed here.** (A) Ghanaian samples have higher *eba181* (P < 0.0001) and *Rh2b* (P < 0.0001) relative expression than Malaysian samples. (B) No significant differences are observed between Ghanian and Gambian samples. Horizontal lines represent the mean relative expression of the sample groups. Mann Whitney test using multiple comparisons has been applied. Only significant differences are shown. Significant values are labelled with * for P < 0.01, with ** for P < 0.001 and with *** for P < 0.0001.

### 3.3. Comparison of *eba* and *Rh* gene transcript proportions with previous data on clinical isolates from another African population

Previously, analyses of the same eight *eba* and *Rh* gene transcripts had been performed on schizont-enriched preparations of other African clinical isolates in the first cycle of *ex vivo* culture, using identical RT-qPCR methods within the same laboratory (Bowyer et al., 2015). The largest available African dataset was from Ghana (N = 26), allowing direct comparison with the Gambian and Malaysian isolates examined in the present study. There were no significant differences in the comparison of data on Gambian isolates from the present study and the previous data from the Ghanaian isolates for any of the gene transcripts (Fig. 3). In contrast, Malaysian isolates displayed higher levels of *eba175*, though not significant statistically (P = 0.0688), and lower levels of *eba181* (P < 0.0001) and *Rh2b* (P < 0.0001) than the Ghanaian isolates (Fig. 3), consistent with differences between the Malaysian and Gambian data (Fig. 1).

Exploratory correlation analyses were performed to examine covariation among ligand transcript levels (Supplementary Fig. S1). Several correlations between pairs of genes were significant in data from one of the populations but not in others, although the direction of correlation was generally consistent. For example, *Rh2b* and *Rh4* expression correlated positively in The Gambia (Spearman’s rho = 0.61, P < 0.004), with weaker non-significant positive correlations in Ghana and Malaysia (rho = 0.37 and 0.38 respectively). The strongest and most consistent association across all populations was the positive correlation between *Rh2a* and *Rh5* transcript levels, (Malaysia, rho = 0.88, P < 0.0001; The Gambia, rho = 0.73 P < 0.001; Ghana, rho = 0.63, P < 0.001) (Supplementary Fig. S1).

## 4. Discussion

This study indicates that, compared to endemic *P. falciparum* populations in Africa, relative levels of different *eba* and *Rh* merozoite ligand genes were more distinct in Malaysia when *P. falciparum* was still endemic there. In particular, *eba181* and *Rh2b* transcript levels were lower, whereas *eba175* levels were higher in Malaysia. Nonetheless, most of the total variation occurred within populations rather than between them.

This is the first comparison of variation of expression in these genes between Asian and African *P. falciparum* populations, and suggests that further studies of Asian parasite populations would be useful to establish whether differences are particular to geographical regions. Studies of transcriptomes in schizonts also indicate other genes with significant variation in expression among different parasite isolates (Hocking et al., 2022; Tarr et al., 2018) although varying methods among studies have not enabled direct comparisons between populations from Africa and Asia. Compared with the achievement of having a global database of genomic sequence variation in *P. falciparum* (MalariaGEN Wellcome Open Research 2025), there are more challenges in building a standardised transcriptome database covering different parasite populations, although benefit may be extended by a single-cell approach (Tripathi et al., 2022).

Variation in transcript levels of different *eba* and *Rh* genes is only one level that might affect functional differences in invasion pathways. It is also possible that other differences affect the abundance or subcellular location of individual ligand proteins, although this is harder to measure. Furthermore, common allelic polymorphisms encode alternative amino acid sequences that may affect ligand binding to erythrocyte receptors (Maier et al., 2009). There were no significant correlations between the relative levels of ligand gene expression and varying parasite multiplication rates among the different isolates, except among the Malaysian samples for *eba181* alone. Further studies on *P. falciparum* clinical isolates in Southeast Asia are needed to determine whether variation in the levels of this ligand have an effect on parasite multiplication rate variation in the region, or whether this correlation appeared by chance.

Moreover, the individual roles of these set of proteins go beyond the alternative routes of invasion. Rh1 and Rh2b have been found to activate calcium signalling inside the merozoite, allowing the release of EBA175 (Aniweh et al., 2016; Gao et al., 2013). Such a stepwise coordination of events, added to translation and protein modifications that can affect the ligand expressions, further illustrate that RNA transcript analysis alone will not correlate with all functional variation. In addition, Rh2b and a fragment of Rh1 have been found collocating with Apical Merozoite Antigen 1 (AMA1) in the tight junction formation (Aniweh et al., 2016; Gunalan et al., 2020), which takes place after the initial binding interactions.

This comparison shows that the Malaysian isolates have significantly different expression profiles compared to either isolates from The Gambia studied here or isolates from Ghana that had previously been assayed. In contrast, there were no significant differences between the African populations, although the Ghanaian isolates had been studied by sampling schizonts in the first *ex vivo* cycle whereas the Gambian isolates were studied by sampling schizonts after a few weeks of continuous *in vitro* culture. This latter comparison supports previous data suggesting that most clinical isolates do not undergo major changes in patterns of ligand gene expression over the initial weeks of continuous culture (Gomez-Escobar et al., 2010). Given that analyses of transcriptional variation can vary between different laboratories, comparisons in the present study were focused on parasite isolates from different sources that were cultured within the same laboratory before application of identical gene-specific RT-qPCR analyses.

It remains to be determined whether the differences in the transcript levels among the populations are primarily due to the geographical differences, or if they are driven by different levels of infection endemicity. In populations with higher endemicity, acquired immune responses are likely to be stronger and parasites may be under immune selection that affects the profile of ligand gene expression. It is possible that higher expression of *eba181* and *Rh2b* and lower expression of *eba175* in African compared to Malaysian isolates could reflect previous immune selection, particularly if EBA175 is a more effective target of naturally acquired responses than EBA181 or Rh2b. These findings should encourage studies in other endemic populations to support future vaccine design, considering both population differences and the wide spectrum of expression profiles within each population.

## Acknowledgements

We are grateful to Khamisah Binti Abdul Kadir for managing the cryopreserved collection of isolates, and arranging the sample documentation and information for sample shipment. We are grateful for the support of many colleagues at the MRC Unit in The Gambia at LSHTM where sampling was conducted in the context of ongoing studies of malaria, including Prof Umberto d’Alessandro, Mrs Haddy Nyang and Mr Simon Correa. We are grateful to patients and community members for their voluntary participation, hospital staff and community field workers, and those managing laboratory facilities and equipment as well as storage and shipment of materials. The study was supported by funding from the UK Medical Research Council (Project grant MR/S009760/1) to DJC. Sampling of community and hospital infections in The Gambia was supported by a joint MRC/LSHTM fellowship for AC, and an MRC Career Development Fellowship MC_EX_MR/K02440X/1 for AAN.

## CRediT authorship contribution statement

Elena Lantero-Escolar: Conceptualization, Formal analysis, Investigation, Methodology, Validation, Visualization, Writing – original draft, Writing – review and editing.

Lindsay B. Stewart: Investigation, Methodology, Project administration, Resources, Validation, Writing – review and editing.

Balbir Singh: Investigation, Writing – review and editing.

Antoine Claessens: Investigation, Writing – review and editing.

Alfred Amambua-Ngwa: Investigation, Writing – review and editing.

David J. Conway: Conceptualization, Data curation, Formal analysis, Funding acquisition, Investigation, Methodology, Project administration, Resources, Supervision, Validation, Writing – review and editing.

---

**Supplementary Figure 1: Correlations between the relative transcript percentages calculated with Spearman’s rho, for Gambian (A) and Malaysian (B) samples. Correlations between invasion ligand transcript proportions analysed from Ghanian (C) samples reported in a previously published study** (Bowyer et al., 2015). Rho values are included in the chart, and they are indicated in red when the correlation is negative and in blue when the correlation is positive. Only positive significant correlations are labelled because some negative correlations are inevitable given it is relative proportions being compared.The positive significant values are labelled with ** for p < 0.01, with *** for p < 0.001 and with **** for p < 0.0001. Analysis in GraphPad PRISM 10.

Supplementary Table 1: (EXCEL file) Sample information from each isolate, grouped by origin: proportions of each invasion ligand transcript as percentage of total, PMR, sample genotypes (single or mixed), and days since culture started until RNA collection from each sample.

