## Supplementary figures and images for "*Plasmodium falciparum* invasion ligand gene transcript profiles in different populations"

### Supplementary Figure 1

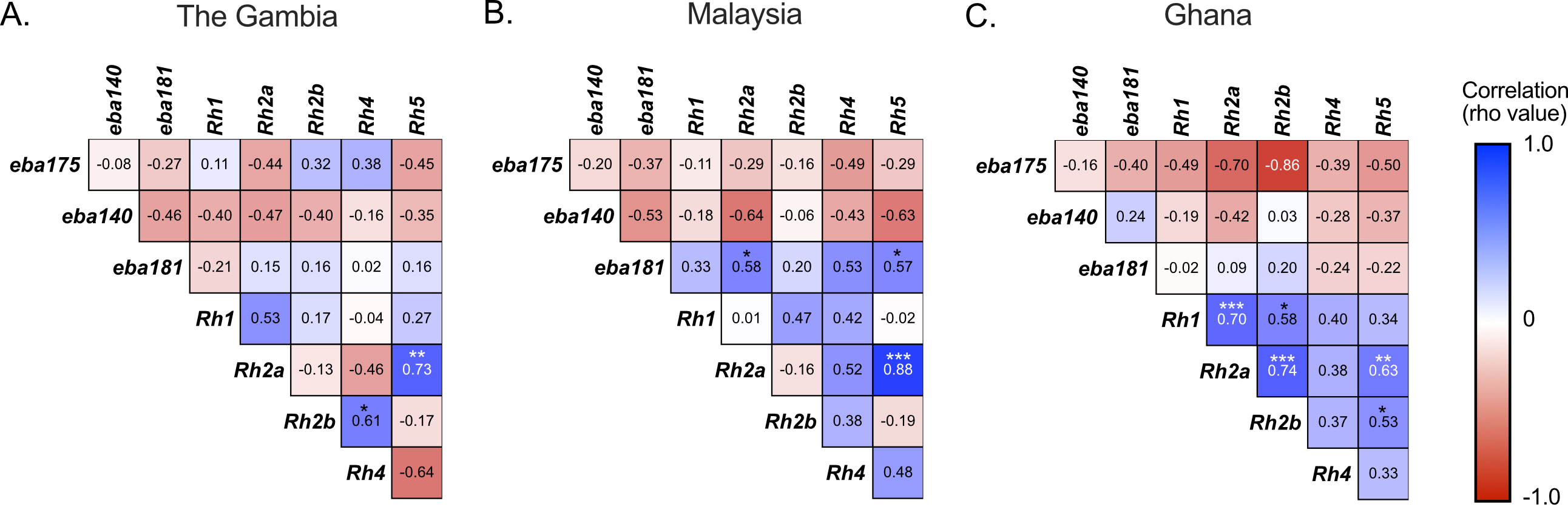
